# Oxygen consumption rate in *Gasterosteus aculeatus-Schistocephalus solidus* system from a non-migratory naturally infected population

**DOI:** 10.1101/111112

**Authors:** Tarallo Andrea, D’Onofrio Giuseppe, Agnisola Claudio

**Affiliations:** Dept. of Biology and Evolution of Marine Organisms, Stazione Zoologica Anton Dohrn, Villa Comunale, 80121 Naples, Italy; Dept. of Biology, University of Naples Federico II, Complesso Universitario di Monte Sant’Angelo, Edificio 7, Via Cinthia, 80126 Naples, Italy

**Keywords:** GASTEROSTEUS ACULEATUS, SCHISTOCEPHALUS SOLIDUS, HOST-PARASITE SYSTEM, CO-EVOLUTION, MIGRATION, LIFESTYLE, RESPIRATION RATE, OXYGEN CONSUMPTION

## Abstract

The three spine stickleback *Gasterosteus aculeatus* is a specific obligatory intermediate host for the cestode worm *Schistocephalus solidus*. This system is commonly used to investigate the host-parasite interaction in fishes. Despite the interesting attempts which have been made to quantify the impact of the parasite over the respiration rate of the host fish, none of the previous reports took in consideration that stickleback is diversified in different ecotypes according to its ability to made reproductive migration, from and to the sea. Here the oxygen consumption rate in specimens of three-spine stickleback collected from a non-migratory population was quantified with the aim to test if the *S. solidus* infection drives a change in the oxygen consumption level of the host fish. The results showed that the infected fishes have a higher rate of oxygen consumption compared with the uninfected one. The differences were due to a direct effect of the parasite, not merely to its contribution to the whole oxygen consumption rate. The data were compared with previous reports, showing that the non-migratory population was characterized by a different level of oxygen consumption rate. The differences were interpreted in terms of divergence in physiological adaptations which had to be appeared in different populations.

## Introduction

Parasitic relationships are very common among living organisms. The *Gasterosteus aculeatus-Schistocephalus solidus* system is a widely used experimental model for investigating the host-parasite interaction in fish (Barber and Scharsack, 2010). The life cycle of the cestode worm *S. solidus* is characterized by three stages. The first starts in the haemocoel of freshwater copepods, which ingest the free swimming embryos of the worm. The *G. aculeatus*, commonly known as three-spine stickleback acquires the cestode larvae alimenting on infectious copepods. *G. aculeatus* is a species-specific host for *S. solidus*. At this stage, the parasite grows up to the adult size within the abdomen cavity of the fish. Sexual maturity is achieved in the third definitive host, which must be an endotherm, normally a piscivorous bird that, by ingesting the infectious stickleback, allow the settlement of *S. solidus* in the intestine. The fertilized eggs are finally released into the water through the bird feces, and a second-generation cycle can occurs. However, the *S. solidus* life cycle can develops in fresh or brackish waters only. Three-spine stickleback, on its side, repeatedly evolved in freshwater-resident ecotypes from ancestral marine/anadromous populations, which could colonize newly inland waters only formed after the retreat of glaciers (Bell and Foster, 1994).

The study of the host-parasite system encompasses the quantification of its energy budget. Many efforts, indeed, have been made to evaluate the influence of *S. solidus* on stickleback energy expenditure (reviewed in Barber et al., 2008), including interesting attempts to determine the whole respiration rate of the infected fishes in comparison with non-infected individuals (Walkey and Meakins, 1970; Lester, 1971). However, virtually no data were available throughout the literature on the whole respiration rate of infected and non-infected sticklebacks from non-migratory populations. However, it has been reported that the level of activity drives the evolution of higher levels of resting oxygen consumption and respiratory surfaces in fishes (Tarallo *et al.*, 2016). Within this frame, the evolution of different levels of respiration rate could results in differential responses to the *S. solidus* infection.

In the following study the oxygen consumption rate in the *G. aculeatus-S. solidus* system was investigated. The respiration rate in specimens of three-spine stickleback collected from a nonmigratory population was measured. Infected and uninfected fishes showed different routine respiration rate, being that of of the former significantly higher than that of the latter.

Comparing our results with those available in the current literature (Walkey and Meakins, 1970; Lester, 1971), the observation that the presence of *S. solidus* drives the increment in the oxygen consumption of *G. aculeatus* was further supported. In both the migratory and the resident ecotypes, independently from the different respiratory adaptations due to their different lifestyles, the same effect is observed.

## Materials and Methods

Adult individuals of three-spine stickleback *Gasterosteus aculeatus* were collected by fish trap in the Nature Reserve of Posta Fibreno (FR, Italy) between October and January. Specimens were maintained in the facilities of the Dept. of Biology of the University of Naples Federico II, and were acclimated for a minimum of 14 days prior to experiments in 50-l aquaria with dechlorinated, filtered and aerated freshwater (20°C, pH 7.0, 10 h:14 h L:D photoperiod). During the acclimation period the animals were fed daily with Chironomus’ larvae, then left to fast for 48 h prior to the experiments.

Rate of routine oxygen consumption, i.e. fish spontaneously activity, were measured using oxygen electrode probe in a closed system. Specimens were weighed, introduced into an insulated respiration chamber (volume of 320ml; constant temperature of 20°C) and left undisturbed to acclimate under a constant air-saturated water flow, from 0.5h up to 1h. Closing time varies with the weight of the measured sample, never exceeding a total fall in oxygen concentration of about 20%. The linear regression of oxygen concentration decreasing over time gives the amount of oxygen consumed by the animal per unit of time. Soon after the oxygen consumption experiment, each specimen was checked for the presence of the parasite *Schistocephalus solidus.*

The procedures described above were approved by the Animal Care Review Board of the University Federico II of Naples.

Data from Walkey and Meakins (1970) regarding routine respiration rate of stickleback specimens, collected in UK areas, were recalculated. According to the same authors, dry weights were converted in wet weight. Data regarding Canadians migratory sticklebacks were recalculated from Lester (1971). Fish length provided by the author were converted to fish weight using the empirical model of Lenght-Weight relationship for *G. aculeatus* available in FishBase (Froese et al., last access February 2017). The above relationship was based on more than thirty thousand observations from different authors. Knowing the worm/fish weight ratio (Lester, 1971), the worm weight were obtained and added to the fish weight. The resulting weight was referred to the whole parasite-host system wet weight.

Statistical analyses were performed using the software R and the VassarStats facilities available from the website (http://www.vassarstats.net/index.html).

## Results and discussion

The weight of the sampled fish ranged from less than one gram up to more than five grams (Table 1). On average, the whole body weight of the infected sticklebacks is slightly higher than that of uninfected fishes (Mann-Whitney test p-value<0.1). This difference could be due to the presence of *Schistocephalus solidus* found in the abdominal cavity of the infected fishes, which increase their total body weight.

**Table 1.** Body mass and specific oxygen consumption rate for infected and uninfected G. aculeatus

|  | Wet Body Weight<br>(g) | Ox Consumption<br>(mlO<sub>2</sub>*h<sup>-1</sup>) | Specific Ox Consumption<br>(mlO<sub>2</sub>*h<sup>-1</sup>*kg<sup>-1</sup>) |
| --- | --- | --- | --- |
| **Uninfected** | 0,91 | 0,6775 | 744,48 |
|  | 1,27 | 0,5780 | 455,08 |
|  | 1,335 | 0,5389 | 403,69 |
|  | 1,464 | 0,5697 | 389,13 |
|  | 2,65 | 0,6435 | 242,84 |
|  | 3,15 | 0,8473 | 268,97 |
| **Infected** | 2,34 | 1,1088 | 473,85 |
|  | 2,6 | 1,1456 | 440,62 |
|  | 2,94 | 0,9628 | 327,47 |
|  | 3,58 | 2,6956 | 752,96 |
|  | 5,37 | 1,5868 | 295,49 |

Regarding the respiration rate, the log-log equation between mass of uninfected fishes and their specific oxygen consumption was O_2_ consumption=-0.79Mass+2.76 (r2=0.89, p-value<0.01; Figure 1). The regression line of the respiration rate from infected fishes showed an increased residual standard deviation, and the corresponding equation was O_2_ consumption=-0.33Mass+2.81 (r2=0.09, p-value N.S.; Figure 1). Infected fishes showed a higher respiration rate than that of the uninfected ones. The statistically significant difference was about 40 mlO_2_^*^h^−1*^Kg^−1^ (One-Way ANCOVA test *p*-value<0.05).

**Figure 1.**
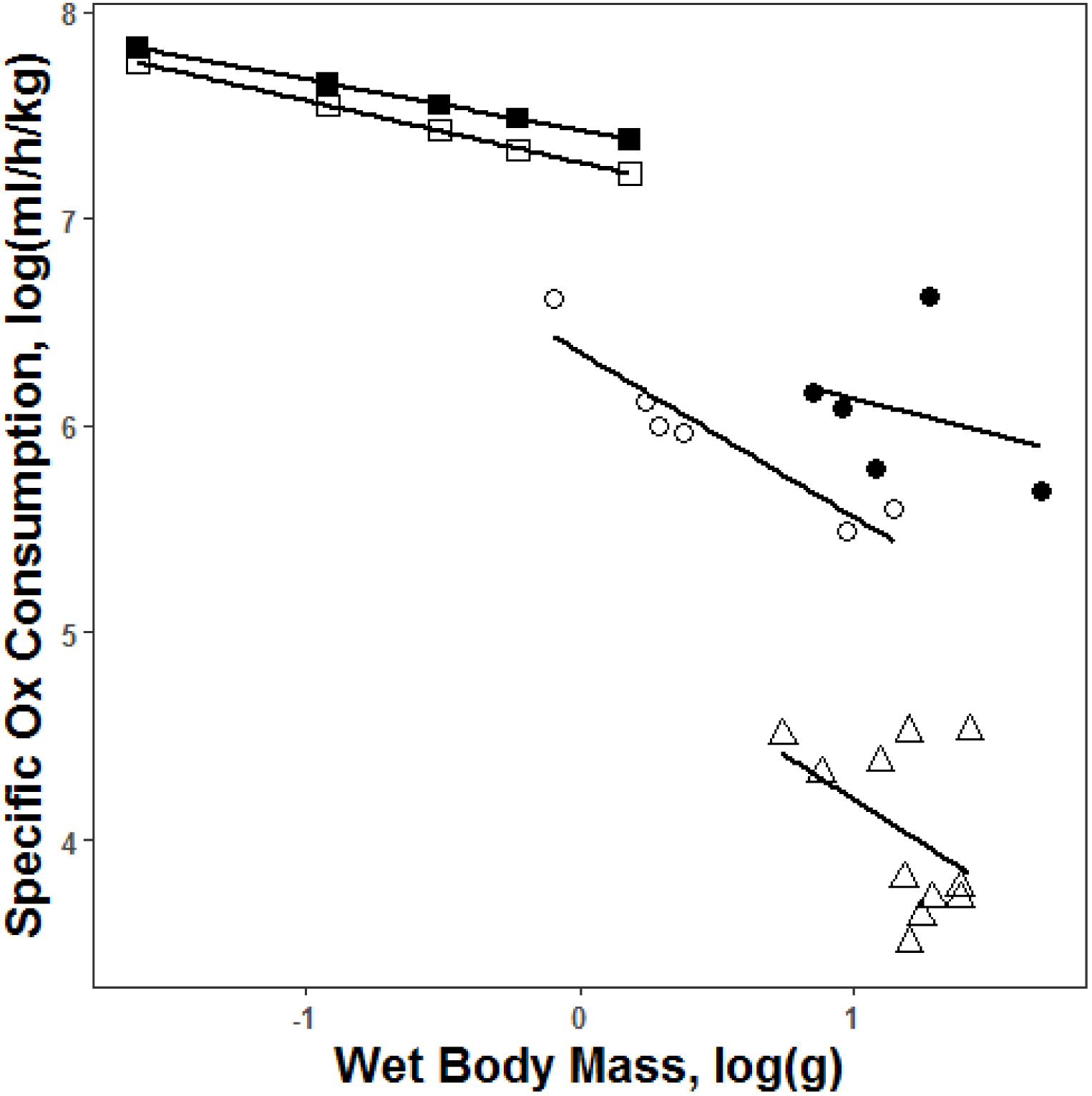
Log-log equation between whole wet body mass versus specific oxygen consumption in infected (full symbols) and uninfected (empty symbols) sticklebacks. UK population are represented by squares (data from Walkey and Meakins 1970); Posta Fibreno lake (Italy) are represented by circles (this study); Aulette lake population by triangles (data from Lester 1971).

The finding was in quite good agreement with previous measurements from Walkey and Meakins (1970). Indeed, the authors reported an intercept of 3.16 and 3.22 for the resting oxygen consumption rate in uninfected and infected *G. aculeatus*, respectively (the equation obtained from their results has been shown in Figure 1), while we observed respectively 2.76 and 2.81. Hence, the difference of intercept between the two regression lines was of the same order of magnitude. Despite that, Walkey and Meakins, contrary to our present results, found no statistically significant difference between infected and uninfected sticklebacks. The authors did not made any statement about the place where the animals sampling was done, though some years later they published a more detailed report on the same topic (Meakins and Walkey, 1975) in which sticklebacks were reported to have been sampled from Eccles, Kent (UK). This, it is not unlikely that the authors repeatedly performed experiments on the same population. Unfortunately, this information could not help to elucidate if the fishes were migrating ecotypes or not, since both the forms are present in that area.

Previous reports highlighted the positive link between level of activity and oxygen uptake in fishes (De Jager and Dekkers, 1974; Tarallo *et al.*, 2016). In particular, migratory sticklebacks are characterized by a standard respiration rate higher than that of non-migratory ones, also when the two ecotypes were sampled from the same river and acclimated at the same laboratory conditions (Tudorache *et al.*, 2007). The authors, extrapolating the standard oxygen consumption from the measured respiration rate during active swimming, revealed an oxygen consumption of 198 ml O_2_^*^h^−1*^Kg^−1^ in the migratory ecotypes, and of 126 ml O_2_^*^h^−1*^Kg in the non-migratory ones (the values are recalculated for sake of comparison with the data here presented). In this frame, the resulting difference between the respiration rate in non-migratory sticklebacks here presented against those produced by Walkey and Meakins (1970) shown a comparable amplitude. The theoretical values for a 1 gram stickleback recalculated from Walkey & Meakins (1970) accounted for 69 ml O_2_^*^h^−1*^Kg. Following the same rationale, the respiration rate for a 1 gram-animal sampled in Posta Fibreno lake accounted for 27 ml O_2_^*^h^−1*^Kg. The difference could be even greater, considering that the temperature at which the experiments were performed by Walkey & Meakins (1970) was lower (15°C) than those for the Posta Fibreno sticklebacks (20°C). Thus, it could be argued that the ecotype of fish measured by from Walkey & Meakins (1970) was the migratory one.

Lester (1971) produced data on migratory ecotype of sticklebacks fished in Alouette Lake (Canada). He found slightly or no differences between infected and uninfected fish (the data obtained from the measures produced for infected fish only were shown in figure 1 for sake of comparison). However, the author pointed out that the mass of the fish carrying the parasite cestode *S. solidus* should be scaled down by the burden of the parasites. Taking this in consideration would increment the estimate of the resting oxygen consumption rate of the host fish alone. Using the equations produced by (Davies and Walkey, 1966) on the respiration rate in *S. solidus*, it can be calculated than a fish should be infected by about 5000 worms 10 grams each to elevate the host-parasite system respiration rate by 1 ml/h. This calculation suggest that the presence of the parasite *S. solidus* drives an increment in the respiration rate of the host through an induced effect, i.e. is not a merely indirect consequence of the presence of the worms consuming more oxygen than the host fish.

Interestingly, infected migratory sticklebacks from Aulette Lake (Canada) (Lester, 1971) have lower oxygen consumption than non-migratory ecotype (Figure 1). The difference was partially due to the lower temperature at which the experiments has been carried on with migratory sticklebacks. The experimental temperature reported by the author was in the range of 10 to 21°C. Also taking this into consideration, the differences were still high. One possible explanation could be that migratory population at Aulette Lake, which cannot migrate anymore because of dams that close the way to the sea, acclimatized by lowering their respiration rate. In seawater fish, often a better growth rate is linked with a lowered surrounding salinity. Usually, this is correlated with a lower standard metabolic rate (Boeuf and Payan, 2001).

As observed by Clarke (1954), the surface of the ponds are preferentially populated by parasitized sticklebacks. This observations has been used as a working hypothesis by Lester (1971) to test if the different distribution of infected and uninfected fish were a consequence of a different respiratory needs between the two categories, i.e. the hypothesis that infected fish tend to swim close to the oxygen rich surface of the pond to satisfy their need for oxygen. Moreover, it has been shown that the higher oxygen consumption was not the only effect of the presence of *S. solidus* in the stickleback abdomen. Barber and coworkers, studying the escape response of experimentally infected *G. aculeatus* to bird predation, reported an altered response of the fish due to the presence of *S. solidus* (Barber et al., 2004). The infected fish showed a slower and sluggish response to fake bird beak. In the wild this behavior should result in an increased risk to be predated by a bird. The net outcome from the *S. solidus* point of view would be the incremented chance to reach the intestine of a third-host bird, where it can complete its life cycle achieving the sexual maturity. No effort has been made to quantitatively clarify if a real segregation of infected fish took place in the water column. As a matter of fact, oxygen concentration is higher at the ponds surface respect to the bottom. The eager of oxygen, typical for parasitized stickleback, should have an influence on their distribution within the water column. It is known that fish are able to sense the oxygen concentration and adjust their behavior to minimize the adverse effects of low oxygen on their fitness (Costa *et al.*, 2014), selecting aquatic oxygen tensions that maintain their metabolic scope for growth and activity (Burleson *et al.*, 2001). In this frame, a lower routine respiration rate of nonmigratory ecotypes could counteract the negative effect of the rise of respiration rate.

In mammals, the elevation of metabolic rate driven by the parasite is well documented, though the exact mechanisms are still unclear (Morand and Harvey, 2000). In ectotherm species exposed to the parasite, an enhancing in the levels of metabolic rate has been reported (Shinagawa *et al.*, 2001; Khokhlova *et al.*, 2002). The energetic costs of maintenance and activation of the immune system probably account, at least in part, for the difference. In fishes, there is no golden rule for respiration rate after parasitic infection. In Arctic charr *Salvelinus alpinus* it has been observed the metabolic depression due to the parasite infection (Seppänen *et al.*, 2008 and references therein). Hence, the host-parasite system *G. aculeatus-S. Solidus*, still a very interesting model, cannot be generalized to other fish-parasite systems.

More effort would be devoted for the unveiling of the full picture, particularly to the study of the physiology of the *G. aculeatus-S. Solidus* system, both in field and in lab controlled experiments.

## Acknowledgements

Authors are grateful to Dr. C. Bailey for challenging discussions that greatly improved the manuscript. AT has been supported by the SZN A. Dohrn research fellowship #04/2016.

## References

Barber, I. and Scharsack, J.P. 2010. The three-spined stickleback-Schistocephalus solidus system: an experimental model for investigating host-parasite interactions in fish. Parasitology, 137: 411–424.

Barber, I., Walker, P. and Svensson, P.A. 2004. Behavioural Responses to Simulated Avian Predation in Female Three Spined Sticklebacks: The Effect of Experimental Schistocephalus Solidus Infections. Behaviour, 141: 1425–1440.

Barber, I., Wright, H., Arnott, S. and Wootton, R. 2008. Growth and energetics in the stickleback-Schistocephalus host-parasite system: a review of experimental infection studies. Behaviour, 145: 647–668.

Boeuf, G. and Payan, P. 2001. How should salinity influence fish growth? Comp. Biochem. Physiol. C. Toxicol. Pharmacol., 130: 411–23.

Burleson, M.L., Wilhelm, D.R. and Smatresk, N.J. 2001. The influence of fish size size on the avoidance of hypoxia and oxygen selection by largemouth bass. J. Fish Biol., 59: 1336–1349.

Clarke, A.S. 1954. Studies on the life cycle of the pseudophyllidean cestode Schistocephalus solidus. Proc. Zool. Soc. London, 124: 257–302.

Costa, K.M., Accorsi-Mendonça, D., Moraes, D.J.A. and Machado, B.H. 2014. Evolution and physiology of neural oxygen sensing. Front. Physiol., 5: 302. Frontiers Media S.A.

Davies, P.S. and Walkey, M. 1966. The effect of body size and temperature upon oxygen consumption of the cestode Schistocephalus solidus (Müller). Comp. Biochem. Physiol., 18: 415–425.

De Jager, S. and Dekkers, W.J. 1974. Relations Between Gill Structure and Activity in Fish. Netherlands J. Zool., 25: 276–308.

Froese, R., Pauly, D. and Editors. n.d. FishBase. World Wide Web Electron. Publ. version (04/2012).

Khokhlova, I.S., Krasnov, B.R., Kam, M., Burdelova, N.I. and Degen, a. a. 2002. Energy cost of ectoparasitism: the flea Xenopsylla ramesis on the desert gerbil Gerbillus dasyurus. J. Zool., 258: 349–354.

Lester, R.J.G. 1971. The influence of Schistocephalus plerocercoids on the respiration of Gasterosteus and a possible resulting effect on the behavior of the fish. Can. J. Zool., 49: 361–366.

Meakins, R.H. and Walkey, M. 1975. The effects of parasitism by the plerocercoid of Schistocephalus solidus Muller 1776 (Pseudophyllidea) on the respiration of the three-spined stickleback Gasterosteus aculeatus L. J. Fish Biol., 7: 817–824. Blackwell Publishing Ltd.

Morand, S. and Harvey, P.H. 2000. Mammalian metabolism, longevity and parasite species richness. Proc. R. Soc. London B, 267: 1999–2003.

Seppänen, E., Kuukka, H., Huuskonen, H. and Piironen, J. 2008. Relationship between standard metabolic rate and parasite-induced cataract of juveniles in three Atlantic salmon stocks. J. Fish Biol., 72: 1659–1674.

Shinagawa, K., Urabe, M. and Nagoshi, M. 2001. Effects of trematode infection on metabolism and activity in a freshwater snail, Semisulcospira libertina. Dis. Aquat. Organ., 45: 141–144.

Tarallo, A., Angelini, C., Sanges, R., Yagi, M., Agnisola, C. and D’Onofrio, G. 2016. On the genome base composition of teleosts: the effect of environment and lifestyle. BMC Genomics, 17: 173. BioMed Central.

Tudorache, C., Blust, R. and De Boeck, G. 2007. Swimming capacity and energetics of migrating and non-migrating morphs of three-spined stickleback Gasterosteus aculeatus L. and their ecological implications. J. Fish Biol., 71: 1448–1456.

Walkey, M. and Meakins, R.H. 1970. An attempt to balance the energy budget of a host-parasite system. J. Fish Biol., 2: 361–372.

